# Chemical bonds in collagen rupture selectively under tensile stress

**DOI:** 10.1101/2022.09.23.509192

**Authors:** James Rowe, Konstantin Röder

## Abstract

Collagen fibres are the main constituent of the extracellular matrix, and fulfil an important role in the structural stability of living multicellular organisms. An open question is how collagen absorbs pulling forces, and if the applied forces are strong enough to break bonds, what mechanisms underlie this process. As experimental studies on this topic are challenging, simulations are an important tool to further our understanding of these mechanisms. Here, we present pulling simulations of collagen triple helices, revealing the molecular mechanisms induced by tensile stress. At lower forces, pulling alters the configuration of proline residues leading to an effective absorption of applied stress. When forces are strong enough to introduce bond ruptures, these are located preferentially in X-position residues. Reduced backbone flexibility, for example through mutations or cross linking, weakens tensile resistance, leading to localised ruptures around these perturbations. In fibre-like segments, a significant overrepresentation of ruptures in proline residues compared to amino acid contents is observed. This study confirms the important role of proline in the structural stability of collagen, and adds detailed insight into the molecular mechanisms underlying this observation.

## Introduction

Collagen is the main constituent of the extracellular matrix (ECM), a very important and functional biomaterial (1–3). Collagen is formed by individual collagen fibrils, which in turn are formed by the entanglement of assemblies of three amino acid strands into a triple helix, so called tropocollagen. Additional interactions between molecules and fibrils through chemical crosslinks stabilise the collagen structure (4). Tropocollagen is characterised by repeating triplets of amino acids, GXY. The glycine residues facilitate the formation of a characteristic triple helix formed by three strands (5). The other two positions, X and Y, are exhibiting more variation, but are enriched in proline (P) and (2S,4R)-hydroxyproline (O), respectively. This enrichment has been linked to structural stability of collagen fibrils (6, 7).

Collagen is providing structural stability to multicellular organisms by absorbing and resisting forces the extracellular matrix experiences. There are two relevant parts to these stability: Firstly, tropocollagen needs to be difficult to deform and break. Secondly, the assembly of the molecules in to fibrils and their assembly into collagen needs to resist mechanical forces as well. It has been noted that the stability of tropocollagen in particular is determined by energetic contributions, i.e. by the strength of chemical bonds and interactions formed, rather than entropic (8).

It is difficult to study these processes at high resolution through experimental techniques, and as a result modelling has been used to gain insight into these processes and determine how mechanical forces are absorbed within collagen. These efforts have led to a description of the force response of tropocollagen (9). After an initial entropic phase (forces of a few pN), unfolding occurs, where the triple helix starts to unwind and lose its helicity. Once this process is completed around 5000 pN, the backbone is stretched, until the tropocollagen ruptures. The unfolding process and the backbone stretch lead to a bilinear extension behaviour (9), with the modelled behaviour closely matching experimental observations (10). Coarse-grained mesoscale modelling allowed similar insight into the force response of cross-linked fibrils (11). The properties of tropocollagen alone are not sufficient to explain the mechanical behaviour of collagen (12). Nonetheless, the deformations of tropocollagen are key to the understanding of these properties, as the molecular stretching and uncoiling are required for the hierarchical mechanics displayed by collagen (12). A detailed insight into these processes can therefore lead towards novel approaches for identifying and treating disease and injury of tissues (13).

Here, we are concerned with two specific parts of the force response of tropocollagen: (i) How tropocollagen responds to weaker forces, where structural changes are introduced, but no bond breaking occurs. This non-reactive regime corresponds to the molecular uncoiling and backbone stretching described previously (9). (ii) When bond breaking occurs, where such ruptures are introduced, i.e. the reactive regime. The aim of this study is to characterise the molecular mechanisms in detail, adding detail to the mesoscopic descriptions provided by others (9, 11).

For the non-reactive regime, it has been proposed that changes to the configurations of proline residues are providing flexibility to the collagen fibrils (14, 15). The five-membered ring characteristic for proline can be in an *endo* or *exo* configuration (5, 16). At equilibrium, around 55 to 60 % of proline residues at the X position are expected to be *endo* (17, 18). There is a small energy barrier between *endo* and *exo*, which have been calculated to be between 0.0 and 0.5 kcal/mol (17), around 0.6 kcal/mol (19), and around 0.3 kcal/mol (18). These calculations match experimental observation of a nanosecond scale transition between the two configurations (20–27). Changes in the relative population between the two configurations are a way to accommodate changes in the backbone configuration, hypothesised to be a mechanism of absorbing forces (14, 15). Indeed, such a change in the *endo/exo* populations is found around Gly to Ala mutations (18). The methyl group in alanine requires more space, pushing the chains apart within the triple helix. As a result a force is exerted on the backbone, and the populations of *endo/exo*-configurations shift. As Y-position hydroxyproline is in the *exo*-configuration, stabilising the collagen fibrils (5, 16), in GPO collagen this mechanism should lead to a shift in *endo/exo*-populations for the X-proline. For GPP collagen, the absence of hydroxyproline should lead to more flexibility, but a similar mechanism should be observable, albeit the response to force should be different from GPO collagen.

Much less is known about the reactive regime, apart from the fact that this process requires unravelling of the triple helix, which may lead to breaking of covalent bonds. While it is not known which bonds are likely to break, experimental work suggest that radical species are formed (28). Radical formation has also been observed in other biomaterials (29, 30). In addition, it has been reported that bond breaking is located in proximity of cross-linking sites (11–13, 28). Our target is to identify potentially preferred bonds to break, and study the effects of mutations and cross-linking on the location of bond ruptures.

The variation in the constituents of native collagen samples complicate studies of collagen. As a result collagen-like models are often used, in particular GPO and GPP repeats that capture some of the important chemical and physical aspects of collagen fibres. Even in these simplified models, it can be difficult to resolve molecular mechanisms in detail. An additional challenge in the context of mechanical behaviour is the application of forces, which complicate experimental setups. If covalent bonds are broken, an additional challenge is the lifetime of potential products, and whether the process and the resulting products can actually be characterised.

As a result, computational methods can support experimental work in this field. For the non-reactive regime, we used the computational potential energy landscape framework (31, 32) to explore the energy landscapes resulting from different forces being applied. The method has previously been used to study the effect of forces on biomolecules (33), and to study the effects of Gly to Ala mutations in collagen (18), showing that the method is suited to resolve force-induced changes in collagen model structures. To study the reactive regime, we implemented a novel simulation scheme to probe the products immediately after the tropocollagen ruptures, employing a semi-empirical potential.

We find evidence for the proposed mechanism of proline ring configuration changes that absorb weaker forces. In the reactive regime, we observe a preference for breaking the C-C*_α_* bond in X-position residues. The introduction of mutations, deletions or cross linking shifts the preference for breaking into the vicinity of these perturbation. In fibre-like segments, the preference for breakages in the C-C*_α_* bond in X-position residues is also observed, with ruptures in proline significantly overrepresented compared to the amino acid content.

## Methodology

In the non-reactive regime, we used the computational energy landscape framework (31, 32) to explore the energy landscapes associated with different forces for GPP and GPO model peptides. In brief, basin-hopping global optimisation (34–36) was used to obtain low-energy starting points for the pulled collagen triple helices. Using discrete pathsampling (37, 38), kinetic transition networks were constructed (39, 40) by locating transition state candidates with the doubly-nudged elastic band algorithm (41–43). Those candidate structures were converged with hybrid eigenvector-following (44), and the respective local minima found by following approximate steepest-descent paths.

For GPO and GPP, the model proteins consisted of seven repeats per strand (21 residues per strand). The end of the strands were capped with methyl groups connected via peptide bonds (CH_3_-CO (ACE) and NH-CH_3_ (NME)). The properly symmetrised (45, 46) AMBER ff14SB force field (47) was used with an implicit Generalised Born solvent representation (48). The forces were applied at the C atom in ACE and the N atom in NME. The applied pulling potential is

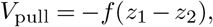

where *f* is the applied force and *z_i_* is the z coordinate of the atom the pulling force is applied to. Even small forces lead to alignment with the z-axis, meaning this setup results in a constant pulling force applied to the molecule. As an AMBER force field is used, the force will be propagated through the harmonic bonding network. The results presented here are for the same force applied to all three strands. The applied forces were 10, 50, 100, 250, 500 and 750 pN.

For simulations in the reactive regime, we first had to establish the forces required to break bonds in these collagen peptides. The potential chosen for this part of the work is the semi-empirical GFN2 within xTB (49), which allows for bond breaking. Harmonic constraints are employed in the terminal residues on either end to ensure that the bond breaking, if it occurs, is located away from the ends of the molecules, introducing more realistic breaking points when comparing the model peptides to tropocollagen. First, we established that bond breaking occurs reliably above forces of 6000 pN by running a series of local optimisations for collagen molecules at various pulling forces. Two energy landscape explorations with xtb were run for forces of 3000 and 4000 pN, as a comparison to the lower force simulations in AMBER. It should be noted that these forces are strong enough to occasionally introduce bond breaking, but not often enough to prohibit the exploration of the energy landscape.

A problem within this reactive framework is posed by the constant force applied. If no fragments are formed, the forces are balanced across the molecules and local minima can be reliably located. However, when fragmentation occurs, the constant force means the fragments will continue drifting apart without convergence to a minimum. One solution is to simply turn of the force at some point, but we considered this as a rather abrupt change in the simulation protocol, which might potentially lead to artefacts in the simulation. Instead, we employed a catching potential. This potential is flat with steep walls, and only yields a significant contribution when the fragments are a specific length *l* apart. This length is chosen to exceed the length of the molecule significantly, so that fragmentation is possible. Once the fragments are separated by *l*, the catching potential counteracts the force, allowing convergence of the gradient and hence the location of local minima. The effect is that the potential catches the fragments and leaves them to hover without the necessity to change the applied force. The catching potential has the form

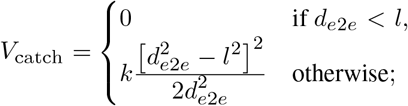

where *d_e2e_* is the end-to-end distance of the molecule, which corresponds to the distance between the atoms to which *V*_pull_ is applied. With this setup, we simulated bond breaking in a number of different sequences, for sequences that had be stretched with a non-reactive force and linked collagen segments. The catching potential allows for rearrangements of the fragments after bond ruptures occurs, and we report here the original fracture site, and not the subsequent changes. For each sequence and condition, a number of local minima where taken from energy landscape databases created for this study or published. Minima were selected at random from these databases. We considered bond breaking in GPO and GPP repeats. In addition, we studied the impact of pre-tensioning and of strand length on the rupture behaviour in GPO repeats. Perturbations to GPO model peptides were introduced in the form of Gly to Ala mutations and Hyp deletions. Finally, more realistic fibre-like segments and cross-linked segments were probed, using collagen models obtained from ColBuilder (50). Table S1 gives an overview of all simulations conducted for this part of the study.

## Results

### Changes in the proline puckering

As the puckering of proline has been suggested as a possible mechanism to absorb mechanical forces, the first property to consider when analysing the simulations for the non-reactive regime is the puckering state of the X and Y positions. The exploration of the energy landscape allows an ensemble calculation of structural properties, including higher energy structures. In Fig. 1 A and B, the puckering configurations are illustrated for GPO and GPP model proteins, respectively. The exact values are provide in the supporting information, Tables S2 and S3. In line with the hypothesis by Chow et al (14), we observe significant puckering changes with increasing force. For GPO, the X-proline residue first flips to an endo state, which near saturation between 250 and 500 pN applied force. At this point, the Y-hydroxyproline starts to flip towards endo states. At higher pulling forces, the Y-hydroxyproline is nearly exclusively in an endo configuration and the X-proline starts to be planar due to the high forces. For GPP, we observe a similar behaviour with more and more endo content. Importantly, the transitions occur at lower forces, as expected from the mechanical stability induced by the hydroxylation of the Y-proline.

**Fig. 1.**
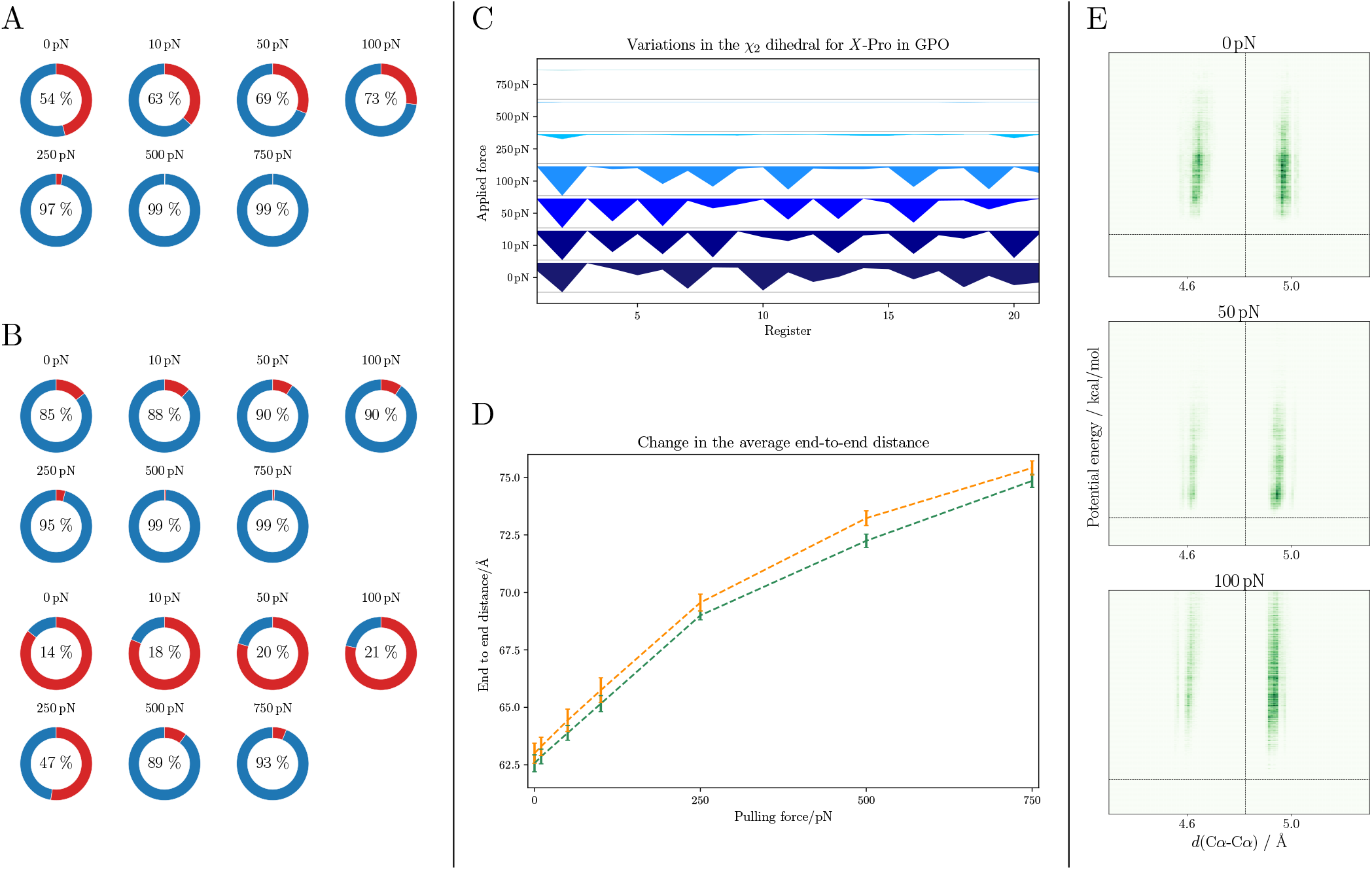
Overview of structural descriptors for non-reactive pulling simulations. **A** Changes to the distribution of X-proline in GPO in *endo* (blue) and *exo* (red) configurations with increasing force. The values are the thermally averaged occupation over the entire molecule (21 registers). **B** Same plots as in A for X-proline in GPP (top) and the Y-proline in GPP (bottom). **C** Variation of the *endo/exo*-distribution across the registers for increasing pulling forces (going bottom to top) in GPO. For each register and force the distribution is shown as the thermally weighted average of occupation for the proline in the register. The *exo*-content is coloured in shades of blue, while the *endo*-content is in white. As the forces increase, the occupation of *exo*-configurations for the X-proline decreases (less and less blue area). Interestingly, the occupation is not uniformly decaying, but instead some residues are nearly exclusively *endo* at low and medium forces. **D** Change in the end to end distance averaged over the three strands and thermally averaged over all structures for GPO (green) and GPP (orange) with increasing pulling force. **E** Heatmaps of the distribution of the distance between the C*_α_* atoms in Pro and Hyp in each register against the potential energy of the minimum. The average distance (18) (dashed, vertical line) and the global energy minimum (dashed, horizontal line) are shown as references. As the force increases from 0 pN (top) to 50 pN (middle) and 100 pN (bottom), the distribution shifts towards a preference for a longer distance, which is corresponding to the *endo*-configuration of X-proline.

The transitions are also not uniform across the registers, in line with findings that the puckering angle is not distributed evenly along GPO repeats (18). Fig. 1 C shows that this effect is pronounced at lower forces, where some registers, which are fairly evenly distributed along the length of the molecule, exhibit high percentages of *exo*-configurations. At higher forces, these disappear rapidly.

This behaviour is also reflected in the end to end distance change observed between 0 and 250 pN pulling force and the change observed between 250 and 750 pN, shown in Fig. 1 D. In the former regime, which corresponds to the subsequent flipping of more and more X-proline rings, we see an elongation of approximately 0.032 Å per pN, while in the latter saturation regime this rate is reduced to around 0.01 Å per pN.

The proline puckering can also be monitored by measuring the distance between the C*α* atoms of the proline and hydroxyproline in the same register. While the regular structure of the GPO model peptide leads to a sharp distribution for the interchain distances for the distance between proline and glycine, and glycine and hydroxyproline, the puckering states of proline lead to two observed distances between proline and hydroxyproline. As the force increases, this distribution more and more tends to a single value, as shown in Fig. 1 E.

### Preferred bond rupture sites in collagen

The first consideration for our study of the bond rupture in collagen models was a survey of different pulling forces to identify the minimum force required to break collagen reliably. For pulling forces around 4000 pN we observed some bond breaking events. For higher pulling forces, we reliably observed such events with a pulling force of 6000 pN being sufficient for consistent rupturing. For even higher forces around 10,000 pN the rupture occurs abruptly, and may introduce artefacts in our simulations. Thus, we used 6000 pN throughout the remaining simulations.

The bond breaking in the GPO model without any pre-stretching showed a clear preference for bond ruptures in proline residues. The bond most likely to break is the C-C*α* bond leading to the formation of an aldehyde radical and a pyrrolidine radical, with 80 to 85 % of all bond ruptures (see Fig. 2 A (left)). We do not observe any significant effects from considering larger GPO strands with regards to the frequency of bond breakages. In the shorter GPO repeats, we observe a preference for a single proline residue in each strand. This observation may be a finite size effect, but we were not able to find a correlation to the puckering state of the structure pulled. In the larger GPO model peptide, we do not observe such a preference, and the main difference in distribution is related to which residues we restraint in our setup, as shown in Fig. 2 B. When a small pre-tension force is applied, no clear preference in which proline the rupture occurs is observed, but otherwise we observe no changes (see Fig. 2 A (centre)). The second and third most likely fracture sites in GPO are the same bond, but in hydroxyproline and glycine, accounting for 5 to 6 % of all ruptures each.

**Fig. 2.**
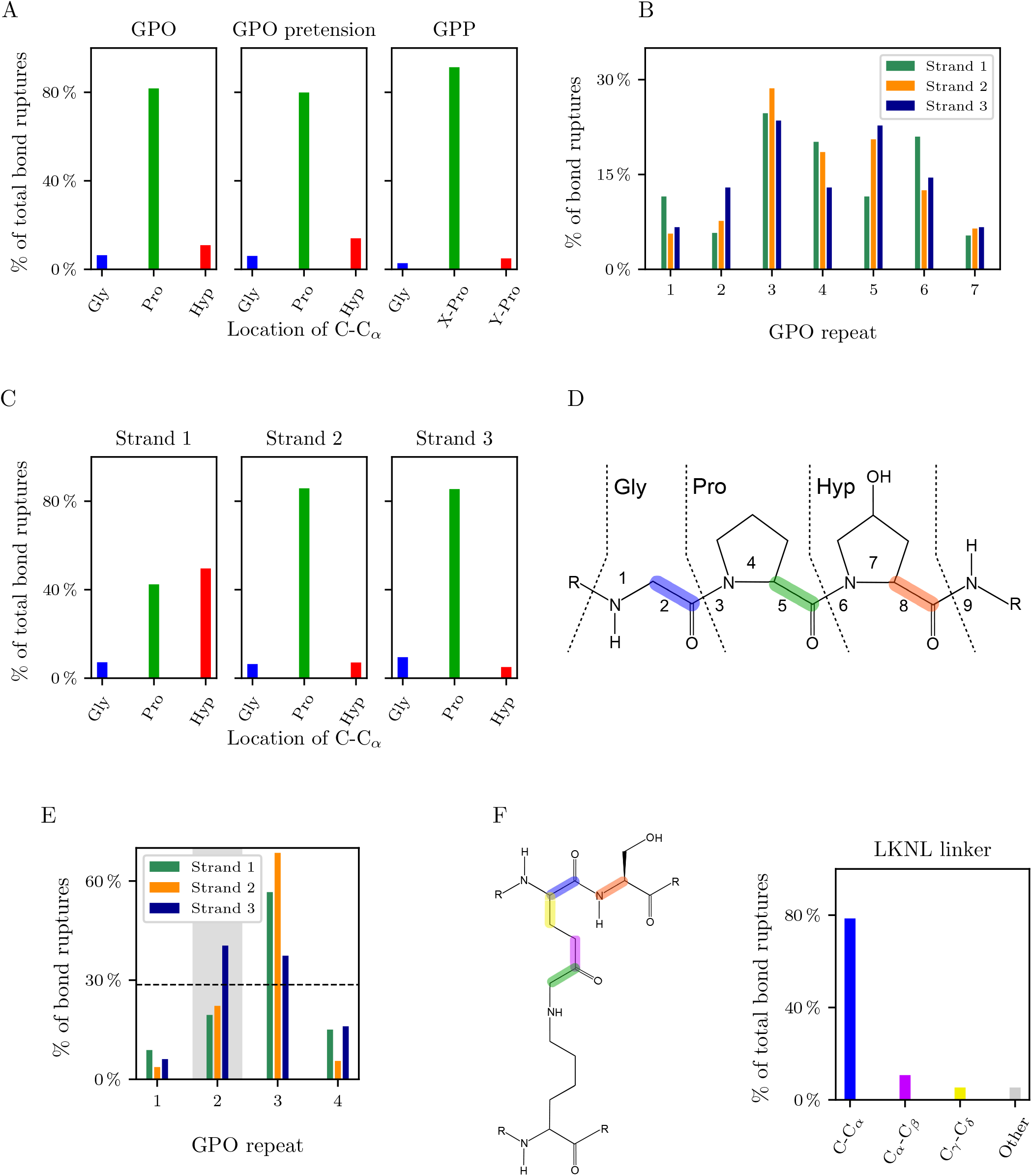
Overview of the results for bond rupture simulations in collagen models. **A** Distribution of ruptures of the C-C*_α_* bond by residue as percentage of the total number of bond ruptures for GPO model peptides (left), GPO model peptides equilibrated at 10 pN (middle) and for GPP model peptides (right) show a strong preference for bond ruptures in the X-position proline. **B** Distribution of ruptures over the seven GPO repeats, where the central 9 registers, which stretch repeats 3 to 6, are unrestrained. There is no clear locality of the ruotures and they are relatively evenly distributed across the sequence, accounting for the constraints applied to the ends of the collagen strands. **C** Shift in breakage location between the mutated strand 1 (one Gly to Ala mutation) and the unmutated strands when a single Gly to Ala mutation is introduced as percentage of the total bond breakages. The significant shift in the left panel, which is the mutated strand, is due to a high likelihood of breaking in the Hyp preceding the mutation. **D** Schematic of the bonds most likely to break as reference for panels A and C. **E** Bond breakage frequencies in interrupted sequences, i.e. where a deletion occurs in repeat 2 in the Y-position in strand 1, leads to ruptures closer to this deletion. **F** Scheme of the bond breaking in the LKNL linker with the relative observed frequency plotted. The highlighted bonds are the only bond for which ruptures were observed, and the percentage of the total ruptures are provided for them. The strong localisation of the bond ruptures around the linker is clear.

In GPP models, we also see bond breaking predominantly of the C-C*α* bond in the X-proline. In fact, it is even more prevalent in GPP, with close to 90 % of all observed bond breakages (see Fig. 2 A (right)).

### Mutations introduce mechanical weakness

In previous work we demonstrated that Gly to Ala mutations lead to a significant change in the backbone around the mutation site, and an associated shift in the puckering state (18). When we apply a rupturing force to these mutated structures, we observe a similar rupture pattern in the unmutated strands as we observe for the pure GPO molecules. However, a significant change occurs in the mutated strands. Here, the rupture occurs in the vicinity of the mutated residue. Nearly 80 % of the ruptures in the mutated strand occur in the two residues on either site of the mutation and the mutated residue, with over 40 % occurring in the preceding residue - in this case hydroxyproline. This results in a significant shift in the rupture location compared to GPO, GPP and the unmutated strands as shown in Fig. 2 D.

### Bond breaking localises around deletions

Another possible change observed in experiment is the deletion of residues in individual repeats. Such interruptions have important biological implications (51–53). Unfortunately, experimental setups are challenging, and generally probe interruptions in all three chain (54), rather than in one strand. Explorations of the energy landscapes for GPO and GPP model peptide-derived interrupted sequences with one deletion show localised effects of the interruption, where kinking of the tropocollagen is observed at the site of the deletion. This kinking has functional importance (51), but also may change the structural stability (53). The hydrogen bonding away from the deletion and between the uninterrupted strands is not effected, while hydrogen bonding involving the atoms in the repeat after the deletion is lost. More detail on these structural details are provided in Supplementary Note 3.

The interrupted sequences show a different selectivity for the rupture locations. While the bond breaking still mostly occurs in X-position proline residues in the C-C_*α*_ bond, the structural alterations due to the deletion impacts where along the strands the bond rupture events occur. The bond breakages occur in the repeats immediately following the deletion, most pronounced in the trailing and middle chain, with over 60 % of ruptures for these strands located in those repeats (see Fig. 2 E).

### Cross-links are mechanically weak

Probing of crosslinks between strands requires the creation of cross linked models. We derived such model using ColBuilder (50), and studied two models for human collagen with dehydro-lysino-norleucine (deH-LNL) and lysino-keto-norleucine (LKNL) linkers. Unfortunately, only one of these models produced results. This finding is related to a subtle change in the simulation setup. Here, we pull the cross-linked strands in opposite directions to load the crosslink. The deH-LNL linker models showed a slight kinking in the strands that are crosslinked. In the simulations this kinking prevented sliding of the chains with respect to each other, and as a result the linker was not loaded. Instead, only the terminal residues in the pulled strands were loaded, and ruptures exclusively occurred in the capping groups. In the LKNL model, the strands were able to slide along each other, allowing for the stress to be distributed along the strands and the linker. This distribution of the force led to breakages in mechanically weak places rather than purely based on the simulation setup. In Fig. 2 F, a scheme is shown for the LKNL linker and the observed bond breaking sites. Apart from a few bond breakages in the residue next to the cross-linked residues, most ruptures occur in the linked residues and the linker. Based on the linking and the setup, one C-C*_α_* bond is loaded and nearly 80 % of the observed bond breaking events are observed in this bond.

### Fibre-like segments exhibit similar behaviour to collagen model peptides

The final set of simulations aimed to identify whether the described patterns so far are representative for more realistic models for collagen, i.e. more complex sequences than GPO and GPP repeats. For this part of the study we tested models for human collagen from ColBuilder (50). The results for the bond breaking in these sequences are shown in Fig. 3. While some variance is observed in which residues contain the rupture location, which is expected from the variance in the amino acid sequence (see Fig. 3), the tendency for breaking of C-C_*α*_ bonds in the X-position residues is still clear. We exclusively see bond breaking in C-C*_α_* bonds, and on average across all strands 78 % of all ruptures are in the X-position residues (see Fig. 3, middle panel), similar to the observations for GPO and GPP repeats (see Fig. 2 A).

**Fig. 3.**
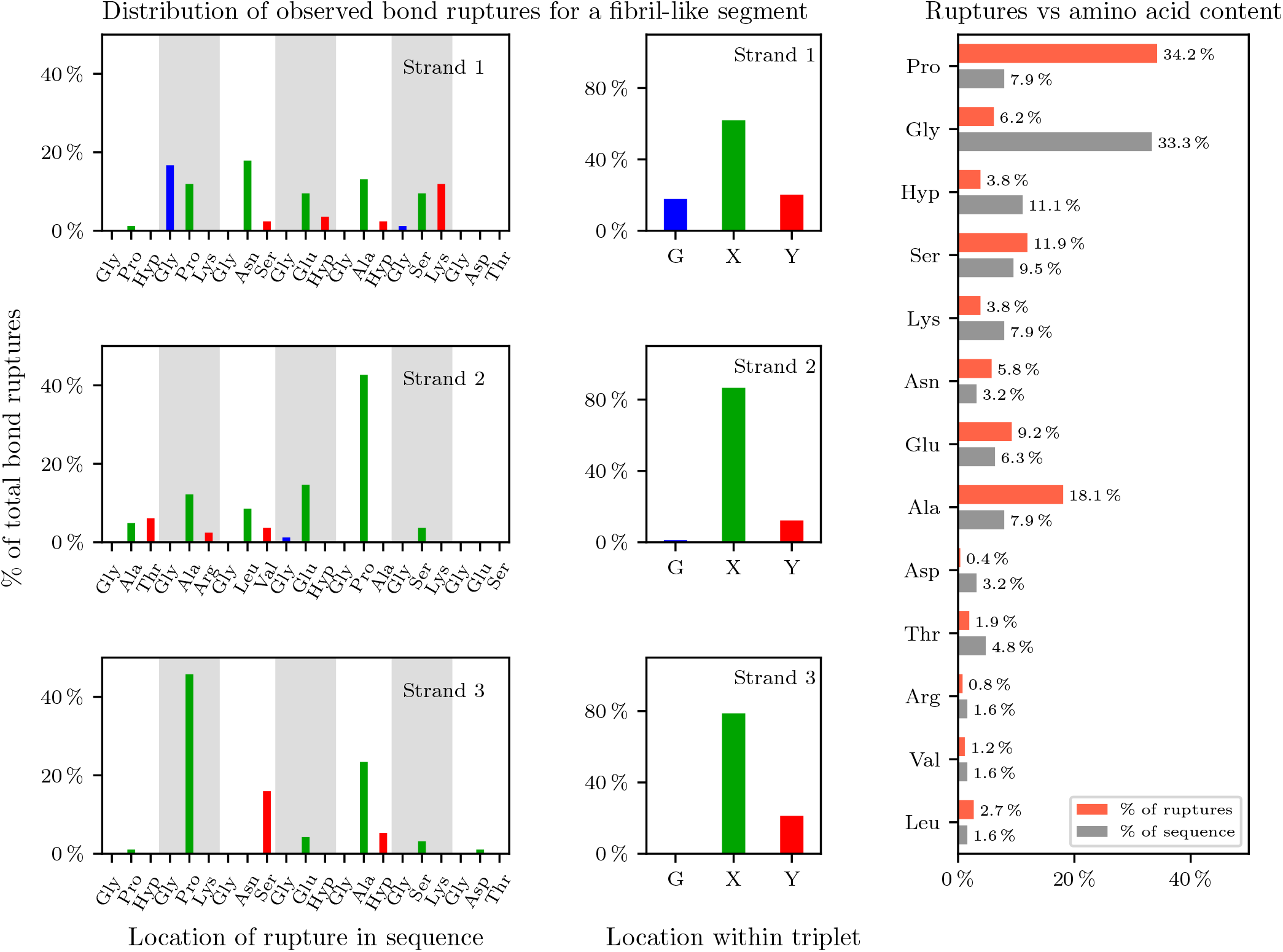
Overview of the results for bond rupture simulations in a fibre-like segment of 21 residues per strand. Left: Distribution of the observed bond ruptures along the sequence. All bonds breaking are C-C*_α_* bonds. Ruptures in Gly are shown in blue, ruptures in X-position residues are green, and those in Y-position residues in red. The individual GXY repeats are highlighted. A preference for X-position ruptures is observed. Middle: Summary of the breakage location by position in the GXY triplets for the individual strands shown in the left panels. Most bond ruptures occur in the X-position (around 78% of all breakages observed). Right: Bond breakages by residue (red) compared to the amino acid content in the fibre-like segment (grey) shows a significant overrepresentation of breakages in Pro compared to its relative amino acid content in the strands. The only other significant overrepresented amino acid is alanine, with glycine and hydroxyproline underrepresented.

As we have more variation in the amino acid content, it is possible to compare the frequency of bond ruptures in specific amino acids to their relative content in the sequence. This information is shown in Fig. 3 in the right panel. The outstanding differences between the two percentages are observed for proline, alanine, glycine and hydroxyproline. In proline residues, 34 % of all bond breaking events are recorded, while only around 8 % of the sequence is made up by proline. A smaller, but still significant increase is observed for alanine, with 18 % of all bond ruptures and only 8 % of the amino acid content. In contrast, both glycine and hydroxyproline show relatively too few bond breaking events. In glycine residues, which make up 33 % of the sequence, only 6 % of the bond ruptures are observed. For hydroxyproline, we observe 4 % of the bond breaks compared to 11 % of the amino acid content.

## Discussion

### Molecular mechanisms of absorption of forces

The first important point to discuss is the mechanism of force absorption observed in the non-reactive regime. All observations in this study confirm the hypothesis of proline *endo/exo* flipping as the mechanism for non-reactive force absorption (14). Importantly, this mechanism extend to ring flipping in Y-position hydroxyproline residues in GPO and proline residues in GPP. The proline ring conformation changes are exhibiting the lowest barrier and occur first.

The changes in the *endo/exo* distribution are not gradual, but flipping seems to occur residue by residue. Once the proline rings are in the *endo* configuration, the end-to-end distance increase is slowed and larger forces are required for the same additional extension. The ring flipping is related to alternatives in the backbone configuration, allowing the backbone to be in better alignment with the helical axis. As a result the end-to-end length for each GPO repeat increases with the applied force. This extension of repeat length is shown in Fig. 4 (left). As a result, GPO repeats, here represented as the vector from the C*_α_* atom in Hyp in one repeat to the same atom in the next repeat, align much better with the helical axis (see Fig. 4 on the right). The proline residues act as flexible extenders, where at no and low applied forces the backbone is winding somewhat more within the framework of the hydrogen bonds that characterise tropocollagen. Larger pulling forces introduce better alignment, and the ring flip allows for this more extended backbone configuration. Likely this process happens segment by segment, as indicated by the residue by residue flipping observed, rather than in a continuous fashion. Importantly, no changes to the hydrogen bonding network is observed at this stage.

**Fig. 4.**
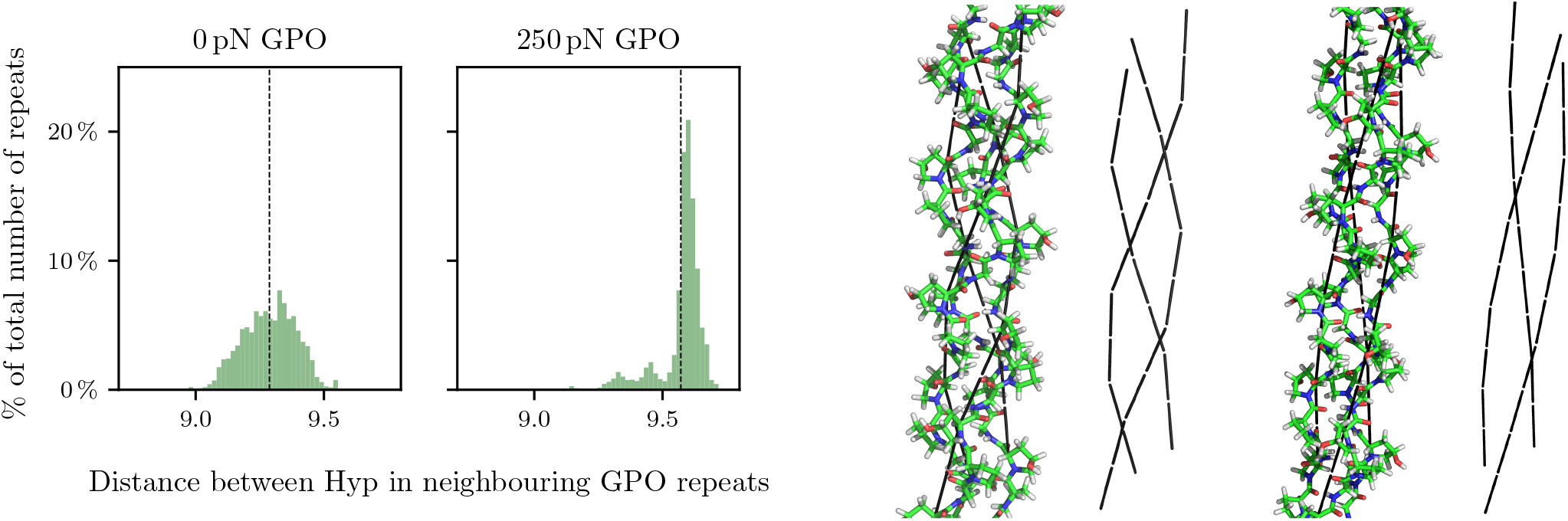
An illustration of the changes in GPO repeat lengths when pulling forces are applied. Left: Distribution of the distance between the C*_α_* atoms in neighbouring hydroxyprolines along the chains as a proxy for GPO length. The average values for the two forces are given by the dashed lines. As the force is increased and proline ring flips are introduced, the segments extend in length and the distribution becomes sharper. Right: The change in force and proline ring configuration leads to changes in the GPO repeat alignment and length. The lowest energy structures are shown for 0 pN (left) and 250 pN (right) with a representation of the vectors between C*_α_* atoms in neighbouring hydroxyprolines shown in black. The forces lead to a higher symmetry and alignment of the GPO repeat orientations.

### Bond breaking in collagen

Once forces exceed the threshold for bond breaking, bond ruptures are readily observed. It should be noted here that not only are our finding for this threshold in broad agreement with macrosocpic models (9), but further that this is significantly larger as the forces required for unfolding observed in non-structural proteins (33). Clearly, this structural resistance to forces stems from the regular hydrogen-bonding pattern and the in-built molecular mechanisms, like the proline ring flipping, and is faithfully reproduced in our simulations.

Two important preferences are observed in the location of the bonds that break. Firstly, the bond most likely to break are C-C*_α_* bonds. This observation holds true throughout all of the models studied. Two reasons can be identified for this bond as the most likely breakage site. Firstly, both resulting radicals, with the exception for glycine, lead to a radical on a secondary carbon and an amidyl radical. Both are somewhat stabilised, albeit rearrangements may occur subsequently. Secondly, radicals are more stable on elements with lower electronegativity, and all other backbone options involve C and N rather than two C atoms.

The second key observation about the location of the bond breaking is the preference for X-position residues. Again This is observed across all models with around 80 % of all observed ruptures in these residues. This preference might stem from the unique position of X-position residues within the collagen assembly. The X-position residue is hydrogen bonded to glycine, and the resulting network is the fundamental contributor to the tropocollagen structure and help to distribute the applied forces throughout the collagen chains. As observed in longer repeats, this distribution works well, will fracture sites fairly evenly distributed across the length of the segments considered. In the GPO and GPP repeats the preference for X-position ruptures automatically leads to a preference of ruptures in proline residues. However, these models are only representative to a point. In this case, we need to consider the fibre-like segments to see whether the same distribution of forces through hydrogen bonding still leads to a preference in ruptures located in proline residues. Indeed, such a preference is observed for these fibre-like models, with a clear preference for ruptures to occur in proline. The mechanism behind this observation is likely related to the force absorption described earlier. Not only to we see ring flipping towards exclusively *endo*-configurations, but moreover at even higher forces rings start to become planar. As a result, the proline bonding is destabilised by the absorption of force, likely leading to the observed strong preference in ruptures.

### The effect of mutations, deletions and cross-linking

The above results yield interesting insight into the molecular mechanisms behind the structural stability of collagen. Additional interest stems from biological changes, such as crosslinking and sequence changes. Crosslinking is key to the formation of strong collagen tissues and crosslinks have previously been identified as a likely place for mechanical ruptures (11–13, 28).

Sequence mutations, such as the Gly to Ala mutation, have been associated with hereditary diseases that impact the mechanical properties of collagen tissues (55). Similarly, while sequence interruptions of the form GX-GXY provide important fucntional binding motifs (51), the same motif in nonbinding motifs may lead to structural destabilisation (53).

For all three of these motifs, we observe localisation of bond ruptures in the proximity of these perturbations. In the cases of the mutation and interruption, the local hydrogen bonding pattern and backbone arrangements are changed. These effects can be seen in the alterations of the ring puckering in the Gly to Ala mutated systems (18) and the kinking and hydrogen-bonding interruption in the interrupted sequences (see Supplementary Note 3). As described above, the hydrogen bonding network distributes forces across the three strands and increases the overall stability. The weakest sites, in the unperturbed models proline residues in X-positions, are then the first to rupture. The interruption of the hydrogen bonding and the restriction of the backbone orientations close to the mutation and deletion sites lead to structural weaknesses, and hence start failing. The crosslinking is not supported by such a hydrogen bonding network, and once the linked chains can slide along each other, the crosslink will be loaded. There is no mechanism to provide additional strength to these parts of the structure, leading to bond breaking in and around the linker, as the collagen strands involved are stabilised.

## Conclusions

In this study we investigated the molecular mechanisms of force response in tropocollagen, considering both nonreactive and reactive responses. For the simulations of reactive responses, i.e. the rupturing of bonds as a result of applied pulling forces, we implemented a novel simulation setup.

For the non-reactive response, we find evidence to support a hypothesis by Chow et al (14) that proline puckering configurations are key to the absorption of applied forces. Not only do we see changes in the X-proline puckering configurations, but also in Y-hydroxyproline and Y-proline in GPO and GPP models, respectively. The X-proline adapts first, before the Y-residues are impacted. Puckering changes seem to happen segment by segment, and lead to longer GXY repeats, which are aligned symmetrically to the helical axis.

When forces are large enough for bonds to break, these breakages are mostly occurring in X-position residues, independent on whether we consider GPO and GPP model peptides or more realistic fibre-like segments. Almost exclusively, the C-C*_α_* bond is ruptured. Rupture sites are fairly evenly distributed in GPO and GPP repeats. When fibre-like segments are considered, we see a strong preference for ruptures in proline residues compared to their relative amino acid content. Likely, this observation is related to the loading of proline residues as force absorbing residues.

When the tropocollagen is altered via mutations or deletions, the interruption of the hydrogen bonding network and backbone orientation changes lead to localisation of bond breakages in the vicinity of these interruptions. Crosslinking similarly leads to localisation of bond ruptures, which matches experimental observations.

Overall, we conclude that bond breaking under tensile stress in collagen is highly selective, and proline residues are central to our molecular understanding of the force response. Mutations and deletions both weaken the collagen chains mechanically, and crosslinks are also weak points within collagen tissues.

## DATA AVAILABILITY

The simulation software (PATHSAMPLE and OPTIM) are publicly available.

The energy landscapes and analysis for the non-reactive regime pulling of GPO and GPP are available on zenodo: https://doi.org/10.5281/zenodo.7107608.

The energy landscapes for the interrupted sequences are also available on zenodo: https://doi.org/10.5281/zenodo.7107558.

The input structures for the reactive pulling simulations are taken from those databases, and the input for the Gly to Ala mutations are taken from previously published data: https://doi.org/10.5281/zenodo.5578060.

The data as well as simulation setup and output for the bond breaking simualtions is provided here: https://doi.org/10.5281/zenodo.7108329. The repository contains the raw count data used for Fig. 2–4 and a summary spreadsheet of all the bond rupture data.

## ACKNOWLEDGEMENTS

The authors thank Prof. Duer for helpful discussions, and Prof. Wales for providing access to computational facilities. KR is funded by a Henslow Research Fellowship from the Cambridge Philosophical Society.

## Supplementary Note 1: Overview of simulations conducted

**Table S1.**
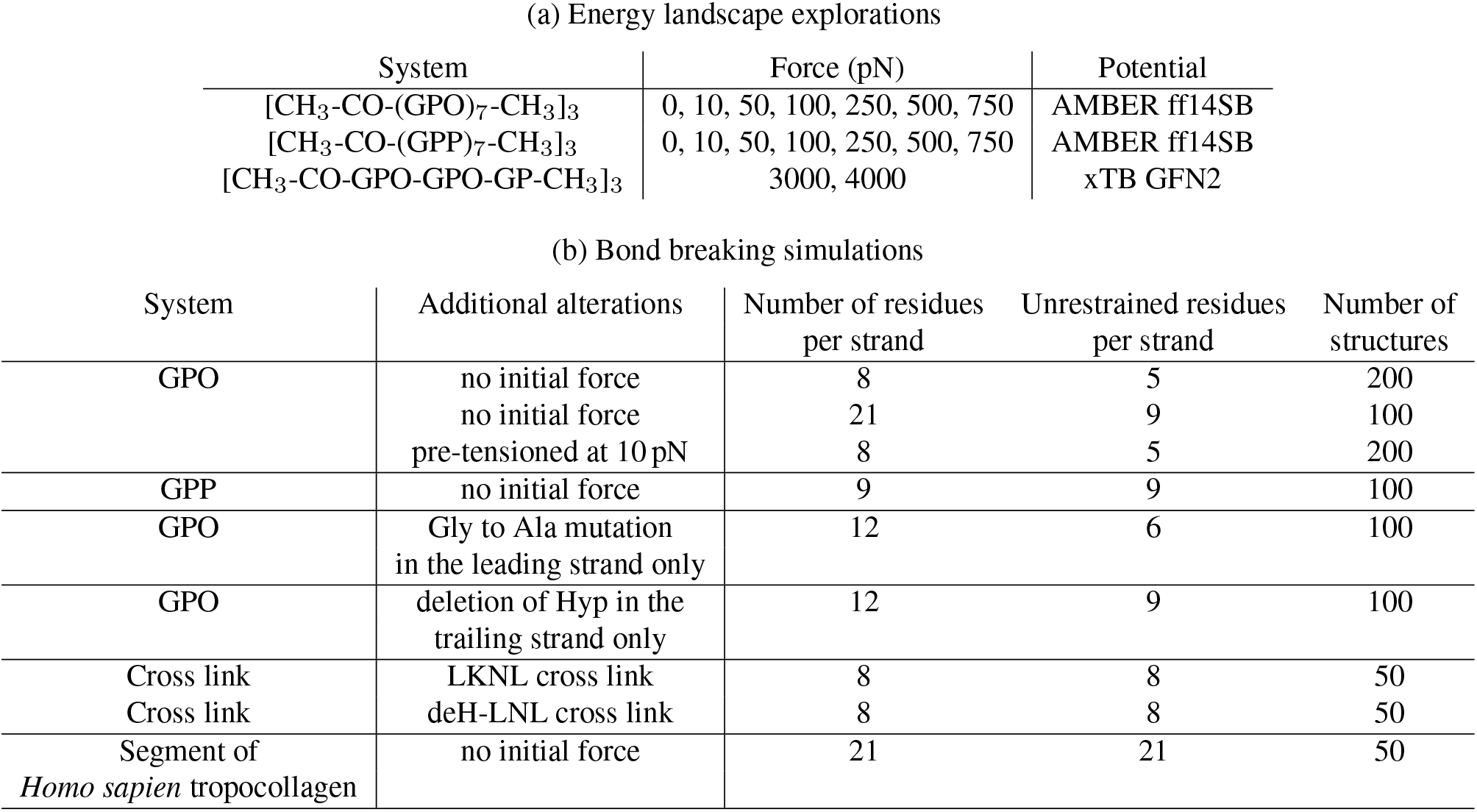
Overview of all simulations conducted for this study. a) Details of energy landscape explorations with DPS, b) details of the bond breaking simulations. All systems contain three chains, apart from the cross-linked systems, which contain six chains.

(a) Energy landscape explorations
| System | Force (pN) | Potential |
| --- | --- | --- |
| [CH<sub>3</sub>-CO-(GPO)<sub>7</sub>-CH<sub>3</sub>]<sub>3</sub> | 0, 10, 50, 100, 250, 500, 750 | AMBER ff14SB |
| [CH<sub>3</sub>-CO-(GPP)<sub>7</sub>-CH<sub>3</sub>]<sub>3</sub> | 0, 10, 50, 100, 250, 500, 750 | AMBER ff14SB |
| [CH<sub>3</sub>-CO-GPO-GPO-GP-CH<sub>3</sub>]<sub>3</sub> | 3000, 4000 | xTB GFN2 |

| System | Additional alterations | Number of residues per strand | Unrestrained residues per strand | Number of structures |
| --- | --- | --- | --- | --- |
| GPO | no initial force | 8 | 5 | 200 |
|  | no initial force | 21 | 9 | 100 |
|  | pre-tensioned at 10 pN | 8 | 5 | 200 |
| GPP | no initial force | 9 | 9 | 100 |
| GPO | Gly to Ala mutation in the leading strand only | 12 | 6 | 100 |
| GPO | deletion of Hyp in the trailing strand only | 12 | 9 | 100 |
| Cross link | LKNL cross link | 8 | 8 | 50 |
| Cross link | deH-LNL cross link | 8 | 8 | 50 |
| Segment of *Homo sapien* tropocollagen | no initial force | 21 | 21 | 50 |

## Supplementary Note 2: *endo/exo* percentages for non-reactive pulling simulations

**Table S2.** Thermal average of puckering percentages for the AMBER (a) and xTB databases (b) for the GPO model protein. For AMBER the endo percentage of the X-proline and the exo percentage for the Y-hydroxyproline are given. Here, missing values to 100% are the exo and endo state, respectively. For xTB, both endo and exo percentages are reported for the X-proline and Y-hydroxyproline. Missing values to 100 % are planar proline rings.

| (a) AMBER databases |  |  |
| --- | --- | --- |
| Forces/ pN | GPO % endo | GPO % exo |
| 0 | 54.05 % | 100.00 % |
| 10 | 63.22 % | 100.00 % |
| 50 | 69.10 % | 100.00 % |
| 100 | 73.05 % | 100.00 % |
| 250 | 97.13 % | 99.99 % |
| 500 | 99.77 % | 99.81 % |
| 750 | 99.96 % | 98.74 % |

**Table S2.** Thermal average of puckering percentages for the AMBER (a) and xTB databases (b) for the GPO model protein.
| (b) xTB databases |  |  |
| --- | --- | --- |
| Forces/ pN | GPO % endo | GPO % exo |
| 3000 | 65.11 % | 2.15 % |
| 4000 | 77.33 % | 0.28 % |
| Forces/ pN | GPO % endo | GPO % exo |
| 3000 | 98.80 % | 0.67 % |
| 4000 | 99.37 % | 0.04 % |

**Table S3.** Thermal average of puckering percentages for the GPP model protein for both the X- and the Y-position proline residues. Missing values to 100% are exo-configurations.

| Forces/ pN | GPP % endo | GPP % endo |
| --- | --- | --- |
| 0 | 85.68 % | 14.47 % |
| 10 | 88.00 % | 18.74 % |
| 50 | 90.79 % | 20.58 % |
| 100 | 90.43 % | 21.61 % |
| 250 | 95.88 % | 47.59 % |
| 500 | 99.28 % | 89.85 % |
| 750 | 99.16 % | 93.81 % |

## Supplementary Note 3: Structural properties of the interrupted sequences

The interrupted sequences consist of GPY repeats, where Y is either O or P, with a single Y-position deleted in the trailing strand. The sequence around that interruption is hence GPO-GP-GPO or GPP-GP-GPP. Energy landscapes were explored for both, GPO and GPP repeats, and the structural ensembles surveyed. In both cases we make two main observation.

Firstly, the structures kink around the deletion, as shown in Fig. S1 (left). The second observation is that all changes are localised around the deletion, as the hydrogen bonding pattern characteristic for tropocollagen is conserved away from the interruption, as schematically shown in Fig. S1 (right).

**Fig. S1.**
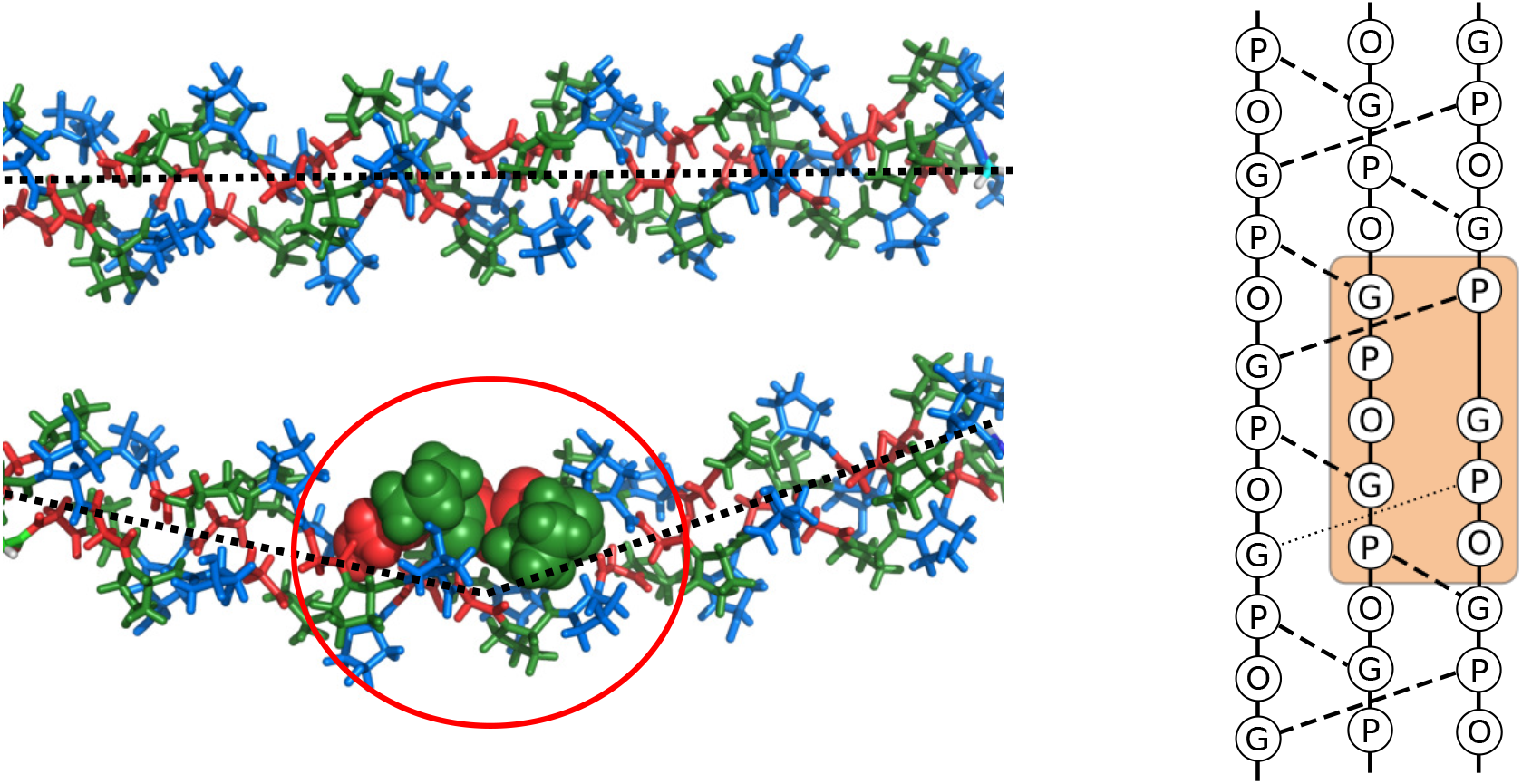
Structural changes observed in GPO repeat sequences when a single Y-position hydroxyproline is deleted in the trailing strand (so called interrupted sequences). Left: Kinking is observed in the interrupted sequence (bottom), around the deletion (spheres), compared to the GPO model peptide straight structure (top). Right: Hydrogen bonding patterns in the collagen with a deletion. The changes are localised around the deletion (box), with the hydrogen bonding between the trailing strand and the other two strands interrupted. The hydrogen bond involving the glycine in the repeat after the deletion is completely absent, and the hydrogen bond involving the proline in the same repeat is partly absent, and only observed in 40% of structures in GPO and 77% in GPP repeats.

